# Metabolic strategies of sharing pioneer bacteria mediating fresh macroalgae breakdown

**DOI:** 10.1101/2021.11.29.470388

**Authors:** Maéva Brunet, Nolwen Le Duff, Tristan Barbeyron, François Thomas

**Author notes:** Address correspondence to François Thomas. The authors have no conflict of interest to declare.

## Abstract

Macroalgae represent huge amounts of biomass worldwide, largely recycled by marine heterotrophic bacteria. We investigated the strategies of “pioneer” bacteria within the flavobacterial genus *Zobellia* to initiate the degradation of fresh brown macroalgae, which has received little attention compared to the degradation of isolated polysaccharides. *Zobellia galactanivorans* Dsij^T^ could use macroalgae as a sole carbon source and extensively degrade algal tissues without requiring physical contact, *via* the secretion of extracellular enzymes. This indicated a sharing behaviour, whereby pioneers release public goods that can fuel other bacteria. Comparisons of eight *Zobellia* strains, and strong transcriptomic shifts in *Z. galactanivorans* cells using fresh macroalgae vs. isolated polysaccharides, revealed potential overlooked traits of pioneer bacteria. Besides brown algal polysaccharide degradation, they notably include stress resistance proteins, type IX secretion system proteins and novel uncharacterized Polysaccharide Utilization Loci. Overall, this work highlights the relevance of studying fresh macroalga degradation to fully understand the niche, metabolism and evolution of pioneer degraders, as well as their cooperative interactions within microbial communities, as key players in macroalgal biomass turnover.

## Introduction

Macroalgae are major primary producers in coastal zones, acting as a global carbon sink (1). Specific polysaccharides dominate macroalgal extracellular matrices (ECM) and can represent up to 50 % of the dry weight (2). For example, brown algae produce alginates and fucos e-containing sulfated polysaccharides (FCSPs). Alginates are linear polymers of β-D-mannuronic (M) and α-L-guluronic acids (G), representing between 10 and 45 % of the algal dry weight (2). FCSPs, accounting for 4-13 % of the dry weight (3), refer to linear or highly branched polysaccharides containing α-linked L-fucose residues together with a variety of other neutral monosaccharides constituents (galactose, mannose, xylose, rhamnose, etc.) and uronic acids (4). They present many substituents, mainly sulfate and acetyl groups. The structure of brown algal polysaccharides is consequently highly heterogeneous and varies according to species, seasons, geographical locations, thallus part, algal growth stages and environmental factors (3-7). Within the ECM, these carbohydrates are cross-linked and associated with proteins (3-15 %), minerals (7-36 % such as iodine, calcium, iron, copper and magnesium), phenols (1-13 %), vitamins, amino acids and small amounts of lipids (1-5 %) to form a complex matrix (8-11). Besides ECM polysaccharides, brown algae also produce laminarin (β-1,3-glucan) and mannitol (12) as storage carbohydrates.

Marine heterotrophic bacteria are crucial for algal biomass mineralization (13). Macroalgae surfaces are constantly colonized by diverse bacterial communities with densities varying from 10^2^ to 10^7^cells cm^-2^ of macroalgal tissue (14). A fraction of these communities, mainly *Bacteroidetes, Gammaproteobacteria, Verrucomicrobia* and *Planctomycetes,* can degrade this complex biomass, showing abilities to hydrolyze purified high molecular weight algal compounds using a considerable enzymatic arsenal (15-18). Over the last 20 years, many studies investigated the algal polysaccharide-processing capabilities of marine heterotrophic bacteria (19), deciphering new catabolic pathways and unraveling the role of carbohydrate active enzymes (CAZymes, http://www.cazy.org, (20)) including glycoside hydrolases (GHs), polysaccharide lyases (PLs) or carbohydrate esterase (CEs), and sulfatases (http://abims.sb-roscoff.fr/sulfatlas/, (21)). In *Bacteroidetes,* CAZymes are usually organized within clusters of coregulated genes involved in carbohydrate binding, hydrolysis and transport, known as Polysaccharide Utilization Loci (PULs). The regulations of these PULs during purified algal substrate degradation were recently studied in a few transcriptome-wide analyses, for both cultivated marine bacteria (22-26) and natural seawater bacterial communities (27). However, using unique substrates does not reflect the complexity of the responses that might occur during the degradation of intact algal biomass. Considering the algae as a whole could reveal novel genes and catabolic pathways, not induced by soluble purified polysaccharides, but playing key roles in algal biomass recycling in the field. To date information on the mechanisms involved in raw algal material assimilation is scarce. “*Bacillus weihaiensis*” Alg07^T^ and *Bacillus* sp. SYR4 grow with kelp and red algal powder, respectively (23,28) and *Microbulbifer* CMC-5 grows with thallus pieces of the red alga *Gracilaria corticata* (29). These studies suggested a successive use of the different brown algal polysaccharides contained in the algal ECM (23) and the release of degradation product in the medium (28,29). However, to our knowledge, no previous work investigated the metabolic mechanisms involved in the degradation of fresh macroalgae, hindering our understanding of algal biomass recycling in coastal habitats. Recently, it has been suggested that among bacteria that are able to use soluble algal compounds, only some populations might be specialists for the breakdown of intact macroalgae tissue (19,30,31). This so-called pioneer bacteria would initiate tissue degradation and expose new substrate niches for less efficient community members considered as scavengers.

The genus *Zobellia (Flavobacteriaceae* family) is frequently found associated with macroalgae and can account for up to 8 % of natural bacterial communities on decaying algae (32-34). It is composed of 15 validly described strains classified in 8 species (35-38). Their genomes encode numerous CAZymes (263-336 genes representing from 6.4 to 7.6 % of the coding sequences), and sulfatases (39-41). Therefore, *Zobellia* spp. are considered as potent algal polysaccharide degraders. In particular, *Zobellia galactanivorans* Dsij^T^, isolated from a red macroalga (35,42), is a model strain to study macroalgal polysaccharide utilization (43). It allowed the discovery of many novel CAZymes and the description of new PULs targeting alginates (44-46), carrageenans (25), agars (47,48), laminarin (49,50), mix-linked glucan (51) and mannitol (52). Its complete transcriptome analysis revealed common regulations triggered by polysaccharides from the same algal phylum (24). *Z. galactanivorans* Dsij^T^ is also well equipped to cope with algal defenses and can accumulate iodine (39,53,54). Moreover, a previous study suggested that *Z. galactanivorans* Dsij^T^ would act as a pioneer bacteria by initiating the breakdown of the kelp *Laminaria digitata,* and demonstrated the crucial role of the alginate lyase AlyA1 in this process (55).

In this study, we aim to better understand the mechanisms controlling fresh macroalgae degradation. To tackle this issue, (i) the complete transcriptome of *Z. galactanivorans* Dsij^T^ was analyzed during the degradation of three brown macroalgae with distinct chemical composition and compared with purified sugars to decipher key genes and mechanisms specifically triggered on fresh tissues and (ii) the ability of *Z. galactanivorans* to degrade fresh algae tissues was compared with other *Zobellia* spp. to assess its singular behavior and hypothesize on potential genetic determinant in fresh macroalgae breakdown.

## Experimental procedure

### Purified substrates

Maltose (Sigma-Aldrich, St. Louis, MO, USA), alginate from *Laminaria digitata* (Danisco [ref. Grindsted FD176], Landerneau, France) and FCSPs from *Ascophyllum nodosum* (Algues & Mer [HMWFSA15424, fraction > 100 kDa], Ouessant, France) were tested for growth. Treatment of this commercial FCSP extract with the alginate lyase AlyA1 (45) followed by Carbohydrate-PAGE (56) revealed it contained alginate impurities. Colorimetric assays (57,58) showed that uronic acids accounted for approximately 24 % (w/w) of the FCSP extract. Based on previous measurements of 9 % uronic acid content in pure FCSPs from *A. nodosum* (59), we therefore estimated the alginate contamination in the FCSP extract to be ca.15%. Alginate, agar (Sigma-Aldrich), kappa-(Goe-mar, St. Malo, France) and iota-carrageenans (Danisco) were used for enzymatic assays.

### Strains

Bacterial strains used in this study are listed in **SuppTable1**, together with previous results of their ability to use pure algal compounds (35-37). They were first grown in Zobell 2216 medium (60) at room temperature before inoculation in marine minimum medium (MMM) complemented with antibiotics to which all the tested *Zobellia* strains are resistant (see supplementary methods for composition) and amended with 4 g.l^-1^ maltose as the sole carbon source. Pre-cultures were centrifuged (3200 g, 10 min) and pellets washed twice in 1X saline solution. Cells were inoculated in microcosms at OD_600_ 0.05.

#### Macroalgae treatment

Healthy *Laminaria digitata, Fucus serratus* and *Ascophyllum nodosum* were collected in May 2019 at the Bloscon site (48°43’29.982” N, 03°58’8.27” W) in Roscoff (France) and cut in pieces (ca. 2.5-3.5 cm^2^) with a sterile scalpel. To clean them from resident epibionts, algal pieces were immersed in 0.1 % Triton in milli-Q water for 10 min followed by 1 % iodine povidone in milli-Q water for 5 min. Finally, algal pieces were rinsed in excess autoclaved seawater for 2 hours, to remove algal exudates and metabolites that could have been produced upon cutting.

#### Microcosm set up and sampling

All experiments were performed in triplicates, except for *F. serratus* in duplicates, at 20 °C in MMM with macroalgae pieces as the sole carbon source. *Z. galactanivorans* was grown in 50 ml with 10 macroalgal pieces, either young *L. digitata* (<20 cm), *F. serratus* or *A. nodosum*. For comparison it was also grown in the same conditions using 4 g.l^-1^ maltose, alginate or FCSPs. During the exponential phase, culture medium (10 ml) and algal pieces were retrieved separately on ice for RNA extraction from the free-living and algae-attached bacteria, respectively. On ice, 0.5 volume of killing buffer (20 mM Tris-HCl pH 7.5, 5 mM MgCl_2_, 20 mM NaN_3_) was added to the liquid samples and cell pellets were frozen in liquid nitrogen. Algal pieces were washed twice in killing buffer:H_2_O (1:1) and frozen in liquid nitrogen. Samples were stored at -80 °C until RNA extraction. To assess *Z. galactanivorans* growth when cultivated in contact or physically separated from algal tissues, incubations were performed in two-compartment vessels (100 ml each) with round bottom and a 65 mm flat edge opening (Witeg [ref. 0861050], Wertheim, Germany), separated by a 0.2 μm filter. Each compartment was filled with 30 ml of MMM and ten *L. digitata* pieces were immersed in one. For comparative physiology, the eight *Zobellia* strains were grown in 10 ml with three *L. digitata* pieces from the meristem part (< 15 cm from the base).

#### RNA extraction and sequencing

Details of the protocols are available in Supplementary Methods. Briefly, free-living bacterial cells were lysed by incubation 5 min at 65 °C in lysis buffer (400 μl) and phenol (500 μl). After phenol-chloroform extraction, RNA was treated 1 h at 37 °C with 2 units of Turbo DNAse (ThermoFisher Scientific, Waltham, MA, USA), purified using NucleoSpin RNA Clean-up (Macherey-Nagel, Hoerdt, France) and eluted in 50 μl of nuclease-free water.

RNA from algae-attached bacteria was extracted as follows. Two algal pieces were immersed in killing buffer, vortexed and placed 7 min in an ultrasonic bath to detach bacteria from the algal surface. Algae were removed and cell pellets resuspended in lysis buffer. RNA extraction and DNAse treatment were performed as described above for the free-living bacteria. To avoid RNA loss on purification columns, DNAse was inactivated using the DNAse inactivation reagent (ThermoFisher Scientific).

DNA contamination was checked by PCR with primers S-D-Bact-0341-b-S-17 and S-D-Bact-0785-a-A-21 targeting the 16S rRNA gene (61). RNA was quantified using the Qubit RNA HS assay kit (ThermoFisher Scientific) and its integrity assessed on a Bioanalyzer 2100 (Agilent Technology, Santa Clara, CA, USA) with the Agilent RNA 6000 Pico kit.

Paired-end RNA sequencing (RNA-seq) was performed by the I2BC platform (UMR9198, CNRS, Gif-sur-Yvette) on a NextSeq instrument (Illumina, San Diego, CA, USA) using the NextSeq 500/550 High Output Kit v2 (75 cycles) after a Ribo-Zero ribosomal RNA depletion step. A total of 24 samples were sequenced (**SuppTable 2**). Sequencing failed for sample Att_Ldig2 due to poor sample quality.

#### RNA-seq analysis

Demultiplexed and adapter-trimmed reads were processed with the Galaxy platform (https://galaxy.sb-roscoff.fr). After read quality filtering using Trimmomatic v0.38.0, transcripts were quantified using the pseudo-mapper Salmon v0.8.2 (62) with the *Z. galactanivorans* Dsij^T^ reference genome (retrieved from MicroScope “zobellia_gal_DsiJT_v2”; Refseq NC_015844.1). Raw counts for individual samples were merged into a single expression matrix for downstream analysis. Raw and processed data were deposited under GEO accession number GSE189322. Principal Component Analysis (PCA) and differential abundance analyses were performed on rlog-transformed data using *DESeq2* v1.26.0 package (63) in R v3.6.2 (64). Genes displaying a log2 fold-change |log2FC| > 2 and a Bonferroni-adjusted P-value < 0.05 were considered to be significantly differentially expressed. The upset plot was done using the *ComplexUpset* package (65,66). Hierarchical clustering was performed using the Ward’s minimum variance method (67). Graphics were prepared using *ggplot2* (68).

#### Enzymatic assays

One volume of 0.2 μm filtered supernatant from the microcosms was incubated with 9 volumes of 0.2 % polysaccharide substrate at 28 °C overnight. Controls were prepared with boiled supernatants. The amount of reducing ends released was quantified using the ferricyanide assay (69). For each sample, the activity measured in controls was subtracted. Finally, the mean value (n=3) measured for the non-inoculated microcosms was subtracted. Significant differences (P<0.05) from 0 were tested using t-tests.

##### CARD-FISH

Algal pieces and culture medium were fixed overnight at 4 °C with 2 % paraformaldehyde. Free-living bacteria were harvested on a 0.2 μm polycarbonate membrane. Catalyzed reporter deposition-fluorescence *in situ* hybridization (CARD-FISH) was performed as described in (34) using the Zobellia-specific probe ZOB137 with helpers. Cells on membrane were visualized with a Leica DMi8 epifluorescent microscope (oil objective 63X). Cells on algal tissues were detected with a Leica TCS SP8 confocal microscope (HC PL APO 63X/1.4 oil objective) using the 488 and 638 nm lasers to detect Alexa488 signal and algal autofluorescence signal, respectively. Z-stack images were collected using 1024x1024 scan format (0.29 μm thick layers, 400 Hz scan speed) and visualized using the surface channel mode of the 3D viewer module (Leica Las X software).

#### Comparative genomics

*Zobellia* genomes were screened for GHs, PLs, CEs and sulfatases using dbCAN2 (70) on the MicroScope platform (https://mage.genoscope.cns.fr). Homologs (>50 % identity and >80 % alignment) were searched for genes of interest using synteny results on MicroScope.

## Results

### *Zobellia galactanivorans* Dsij^T^ degrades and utilizes fresh brown macroalgae tissues as carbon source

*Z. galactanivorans* growth was tested with three brown macroalgae from two different orders and with distinct chemical composition, *Laminaria digitata* (order Laminariales), *Fucus serratus* and *Ascophyllum nodosum* (order Fucales), as the sole carbon and energy source. Growth was detected with the three algal species (OD ≈ 0.2-0.5, **Figure 1A**), with tissue bleaching and damages only visible on *L. digitata* pieces after 65 h (**Figure 1B**). *Zobellia-* specific CARD-FISH assays revealed that even if antibiotic-resistant resident epibionts grew in the non-inoculated controls containing *A. nodosum* and *F. serratus* (one replicate), most of the bacterial biomass after 65h in the Zobellia-inoculated microcosms was *Zobellia* cells (>50 %, **SuppFigure 1**).

**Figure 1.**
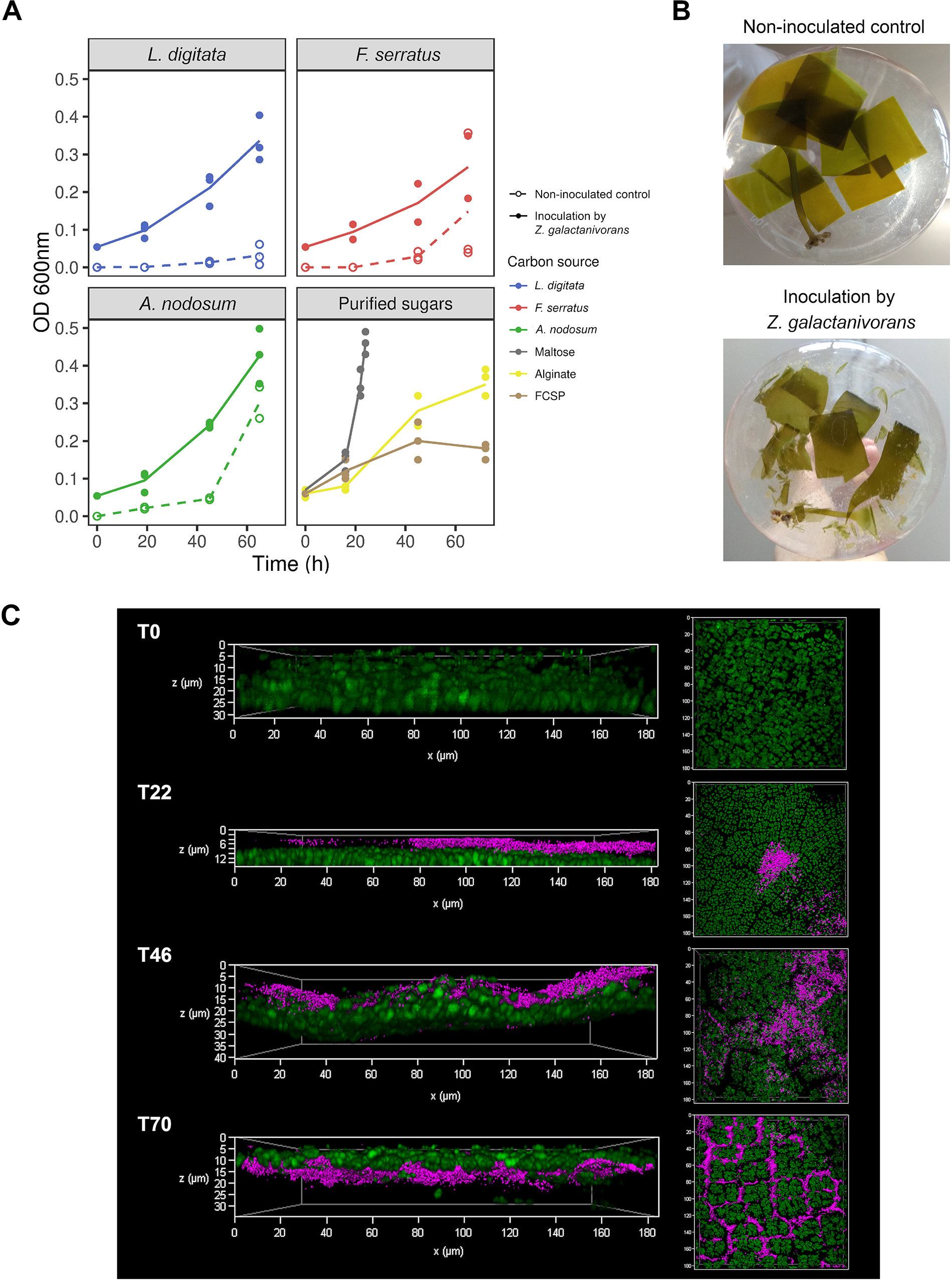
Ability of *Z. galactanivorans* Dsij^T^ to use fresh brown macroalgae for its growth. (A) Growth of *Z. galactanivorans* with either macroalgae pieces (*Laminaria digitata, Fucus serratus* and *Ascophyllum nodosum*) or purified sugars (maltose, alginate and FCSPs). Individual points for replicate experiments are shown. Lines are means of independent replicates (n = 2 or n = 3). (B) Photographs showing the integrity of the *L. digitata* tissues after 65 h. (C) *L. digitata* tissues colonization by *Z. galactanivorans* during the degradation. Micrographs are overlay of the CARD-FISH signal (magenta, Zobellia-specific probe with Alexa488 as the reporter signal) and the algal autofluorescence (green) and were obtained with the surface channel mode of the 3D viewer. For the different times, transversal views are shown on the left and top views on the right. The non-fluorescent gap between the bacterial cells and the algal cells likely represent the mucilage coat of *L. digitata.* The absence of algal autofluorescence signal below 25-30 μm is the result of its rapid decrease in intensity as we move away from the coverslip.

CARD-FISH assays on *L. digitata* tissues showed gradual tissue colonization by *Z. galactanivorans,* from cell patches at the surface of the *L. digitata* mucilage coat to deeper penetration within the tissue invading the intercellular space (**Figure 1C**).

### Transcriptomic shift during fresh macroalgae degradation

*Z. galactanivorans* Dsij^T^ transcriptome of free-living cells obtained during macroalgal degradation was compared to the responses occurring with a disaccharide, maltose, and with purified brown algal polysaccharides, alginate and FCSPs. Between 44 and 93 % of the sequenced reads from free-living bacteria grown with macroalgae mapped on the genome of *Z. galactanivorans* Dsij^T^ (**SuppTable 2**). Multivariate analysis separated samples according to carbon source (**Figure 2A**). Transcriptomes of cells grown with *L. digitata* were closer to that obtained with alginate of FCSPs compared to *A. nodosum* or *F. serratus.*

**Figure 2.**
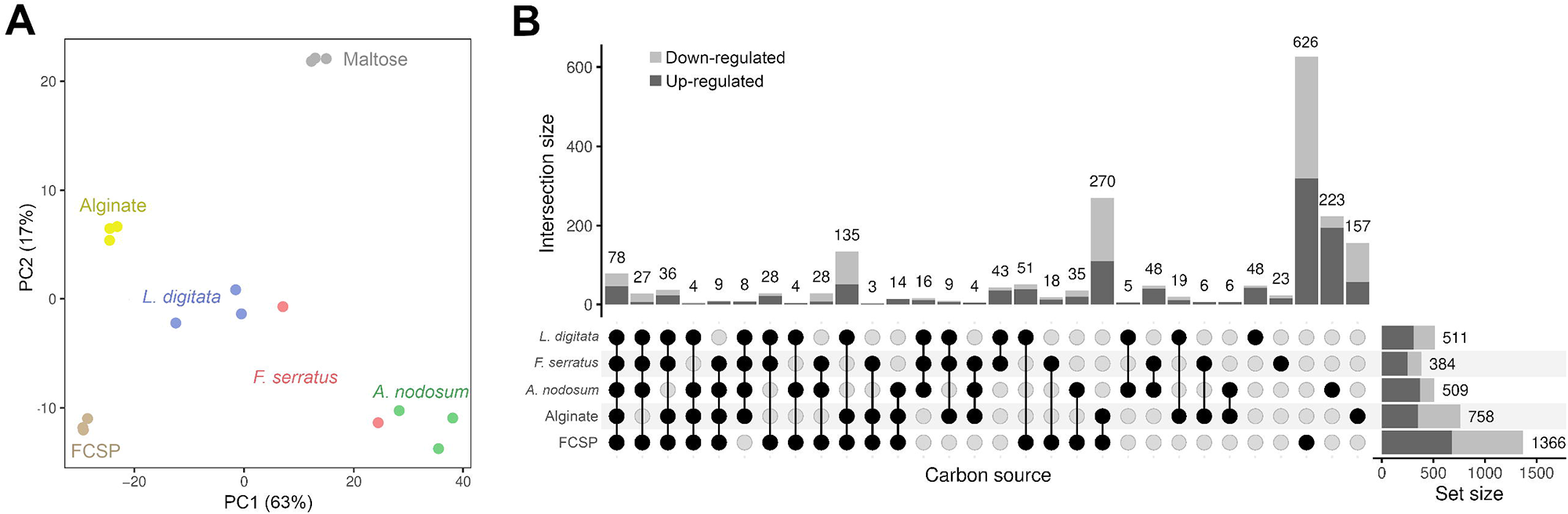
General features of the transcriptomic responses occurring in free-living *Z. galactanivorans* Dsij^T^ during growth with macroalgae. (A) Principal Component Analysis of the gene expression. (B) Upset plot of the differentially expressed genes with maltose as the control condition (Bonferroni-adjusted p-value < 0.05 and |log2FC| > 2). Set size represents the total amount of genes regulated in each condition.

Differential abundance analysis revealed 1117 and 864 genes up- and down-regulated with at least one substrate, using maltose as control (**SuppTable 3**). Among them, 56 % (628 up-regulated genes) and 52 % (449 down-regulated genes) showed substrate-specific regulations (**Figure 2B**). In particular, half of the genes regulated with *A. nodosum* and FCSPs were not differentially expressed in any other conditions. *L. digitata* was the algae inducing the highest number of regulations shared with at least one polysaccharide (399, 254 and 217 genes with *L. digitata, F. serratus* and *A. nodosum* respectively). More regulations were shared between *L. digitata* and *F. serratus* (116 genes) than *F. serratus* and *A. nodosum* (89 genes) or *L. digitata* and *A. nodosum* (13 genes). Finally, a core set of 70 up-regulated and 59 down-regulated genes responded to the three macroalgae.

### Carbohydrate catabolism-related genes

Hierarchical clustering of expression data of the 51 identified PULs in the *Z. galactanivorans* Dsij^T^ genome revealed that PULs predicted to target brown algal polysaccharides grouped together (**Figure 3A**) and were significantly induced with macroalgae. In particular, the alginate-specific PUL29 was significantly overexpressed in all conditions compared with maltose (mean log2FC of 4) and the highest expression was observed with *L. digitata* (**Figure 3B, SuppFigure 2**). Some PULs were exclusively triggered by macroalgae: PUL34 and 35, likely targeting FCSPs (as they encode sulfatases and fucosidases), were significantly triggered by *L. digitata,* PUL4 targeting β-glucan responded to *A. nodosum* and the FCSP PUL3 was induced by both *L. digitata* and *F. serratus.* PUL26 and 27, whose function remains unclear, were both induced by *L. digitata* and FCSPs, as well as by alginate for PUL26 and *F. serratus* for PUL27. Purified FCSPs also induced the expression of 14 PULs outside the described cluster, encompassing a large diversity of targeted substrate (notably β- and α-glucan, sulfated polysaccharides, xylan, unclear substrate). No PUL known to target red algal polysaccharides (e.g. PUL40, 42, 49 or 51) clustered with this set of overexpressed PULs, suggesting a specific induction of brown algal polysaccharide degradation mechanisms in the presence of brown algal tissues. The measured activity of secreted polysaccharidases corroborates this observation (**Figure 3C**), as only the alginolytic activity was significantly higher when *Z. galactanivorans* was grown on macroalgae compared with the non-inoculated control (t-test, P<0.05).

**Figure 3.**
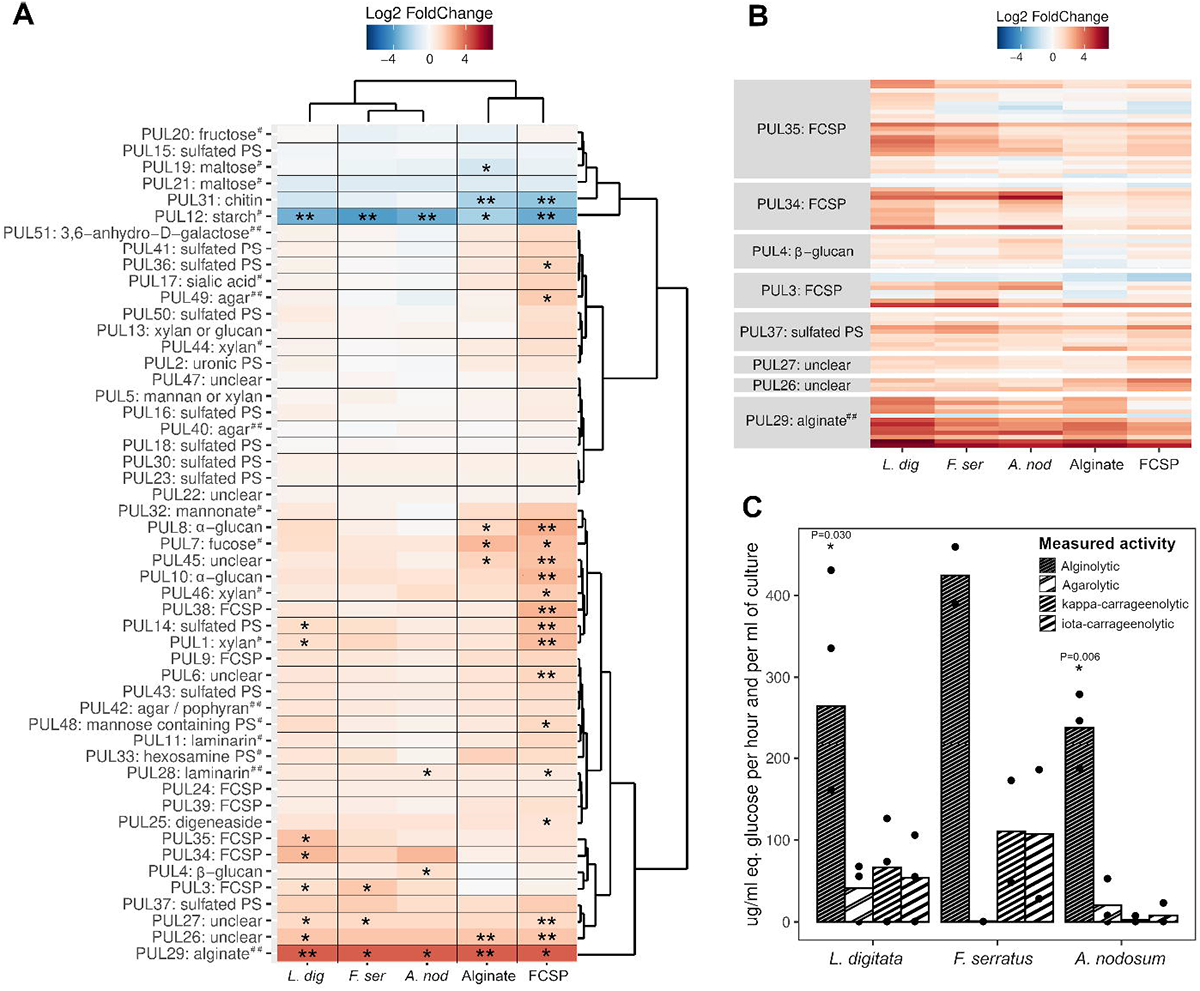
Regulation of the catabolic pathways during the degradation of fresh algal tissues. (A) Heatmap of the 51 PULs identified in the genome of *Z. galactanivorans* Dsij^T^. PUL 1 to 50 were identified during the annotation of the *Z. galactanivorans* Dsij genome by the presence of the susCD-like pair (Supplementary Table S3 in (39)). PUL51 targeting 3,6-anhydro-D-galactose and involved in carrageenan catabolism (but lacking the susCD-like pair) was further described (25). For each PUL, the mean log2FC of all genes is represented, taking maltose as a control condition. Carbon sources and PULs were arranged according to a hierarchical clustering analysis (Ward’s method). A PUL was considered regulated (induced in red, repressed in blue) if more than 50 % of the genes were significantly differentially expressed (*) and strongly regulated if more than 80 % of the genes were significantly differentially expressed (**). Putative substrates targeted by the PULs are indicated. Hash signs denote PULs biochemically characterized previously in *Z. galactanivorans* (##) or in another organism (#). (B) Heatmap representing the log2FC of individual genes contained in the PULs induced with macroalgae and which clustered together in A. (C) Activity of extracellular polysaccharidases secreted in the microcosms containing macroalgae. The mean value measured in the uninoculated controls was subtracted from each value. Bars are means of independent replicates (n = 2 or 3) shown as individual points. Significant difference from zero was tested when n = 3 (t-test; *, P<0.05). *L. dig: Laminaria digitata; F. ser: Fucus serratus; A. nod: Ascophyllum nodosum;* FCSP: fucose containing sulfated polysaccharide; PS: Polysaccharide.

On the other hand, PULs targeting simple sugars (maltose and fructose) or polysaccharides absent from brown algae (starch and chitin) were repressed with macroalgae and purified polysaccharides (**Figure 3A**). The starch PUL12 was strongly underexpressed in all conditions while the chitin PUL31 showed a significant repression only with algal polysaccharides.

### Specific induction with fresh algal tissues

To unravel pathways specifically governing the degradation of fresh macroalgal biomass, we further focused on genes upregulated with at least one macroalgal species compared to maltose and purified polysaccharides. We detected 41, 59 and 189 genes following this pattern with *L. digitata, F. serratus* or *A. nodosum,* respectively (**SuppTable 4**). It included few CAZyme-encoding genes (**Figure 4**), notably two genes within putative FCSP PULs (*zgal_205* [GH117 in PUL3] and *zgal_3445* [GH88 in PUL34]). Other polysaccharidase genes outside classical PUL structures were induced with *A. nodosum,* such as *alyAl (zgal_1182,* alginate lyase PL7), *cgaA (zgal_3886,* glucan 1,4-α-glucosidase GH15), *agaC (zgal_4267,* β-agarase GH16), *pelA1 (zgal_3770*, pectate lyase PL1) and *dssA (zgal_3183*, sheath polysaccharide lyase PL9). GT2 *(zgal_2991, 4154)* and GT4 *(zgal_2990, 3759)* were also triggered with macroalgae. Additionally, many genes linked to oxidative stress responses and Type IX secretion systems (T9SS) were specifically induced with macroalgae (**Figure 4**). A large gene cluster *(zgal_1071-1105)* notably encoding three oxidoreductases, a DNA topoisomerase and a peroxiredoxin was up-regulated with *L. digitata* and *F. serratus.* Other genes encoding antioxidant proteins were triggered, especially on *L. digitata,* such as the superoxide dismutase SodC (ZGAL_114) or a β-carotene hydroxylase (ZGAL_2972), as well as a carboxymuconolactone decarboxylase family protein (ZGAL_1598) which includes enzyme involved in antioxidant defense (71). Two catalases (ZGAL_1427 and ZGAL_3559) were induced in the presence of *L. digitata* and *F. serratus* in comparison to maltose and alginate (**SuppTable4**). Several genes predicted to encode T9SS components were significantly induced during macroalga degradation, in particular with *A. nodosum* (14 out of 33 genes identified in the genome, against 1 and 5 with *L. digitata* and *F. serratus* respectively) (**Figure 4**). They include particularly genes encoding SprF family proteins and T9SS-associated PG1058-like proteins. In addition, 7 unknown proteins containing a conserved C-terminal domain (CTD) from families TIGR04131 (gliding motility – ZGAL_2022, 2761, 2762, 3727) and TIGR04183 (Por secretion system – ZGAL_93, 1124, 4310) were triggered. These CTDs are typical of cargo proteins secreted by the T9SS.

**Figure 4.**
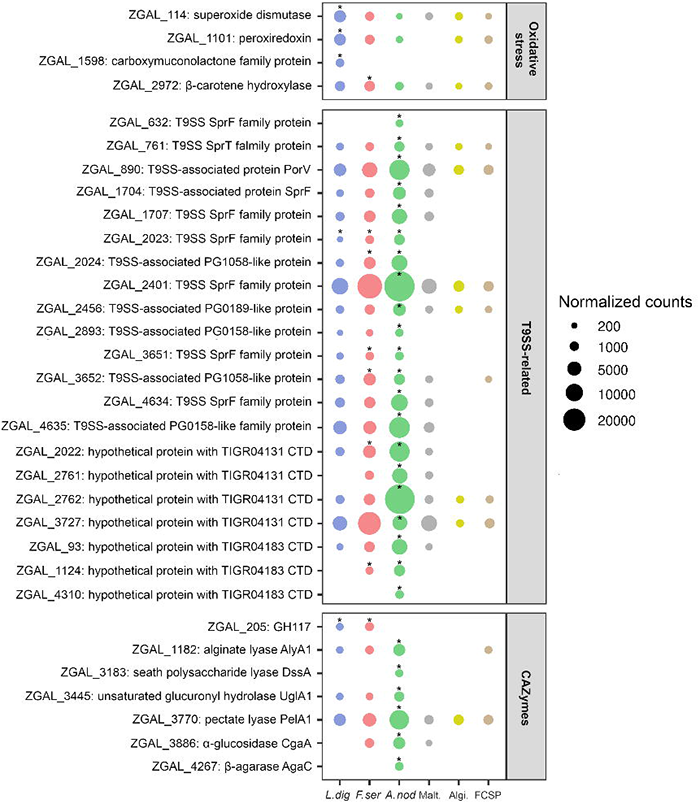
Selection of genes specifically induced by fresh macroalgae. Mean expression values (n = 3, except for *F. serratus* n = 2) of selected genes significantly triggered (*) with at least one macroalgae compared to both the purified polysaccharides and maltose (see SuppTable 4). No dot was represented if the mean read count was below 200. *L. dig: Laminaria digitata; F. ser: Fucus serratus; A. nod: Ascophyllum nodosum;* Malt.: Maltose; Algi.: Alginate; FCSP: fucose containing sulfated polysaccharide

### Effect of bacterial attachment to macroalgae

Transcriptomes of algae-attached cells were compared to that of free-living bacteria. To minimize bias due to poor sequencing depth (caused by a high proportion of eukaryotic rRNA in algae-attached samples [**SuppTable 2**]), we discarded samples from *A. nodosum* microcosms (up to 70 % of uncovered coding regions) and only considered upregulated genes (not the down-regulated ones) in attached versus free-living bacteria. Respectively 19 and 14 genes were significantly induced in bacteria attached to *L. digitata* or *F. serratus* (**Figure 5A**), including a shared set of 5 genes from a genomic region *(zgal_4237-4246)* putatively involved in stress responses. Attachment to *L. digitata* induced the expression of two TetR-type transcriptional regulators, 2 YceI family proteins which might be involved in oxidative stress response via isoprenoide synthesis (72,73) and 5 chaperones (**Figure 5A**). Cells attached to *F. serratus* notably overexpressed 2 TonB-dependent receptors not associated with a SusD-like protein, likely not involved in carbohydrate metabolism.

**Figure 5.**
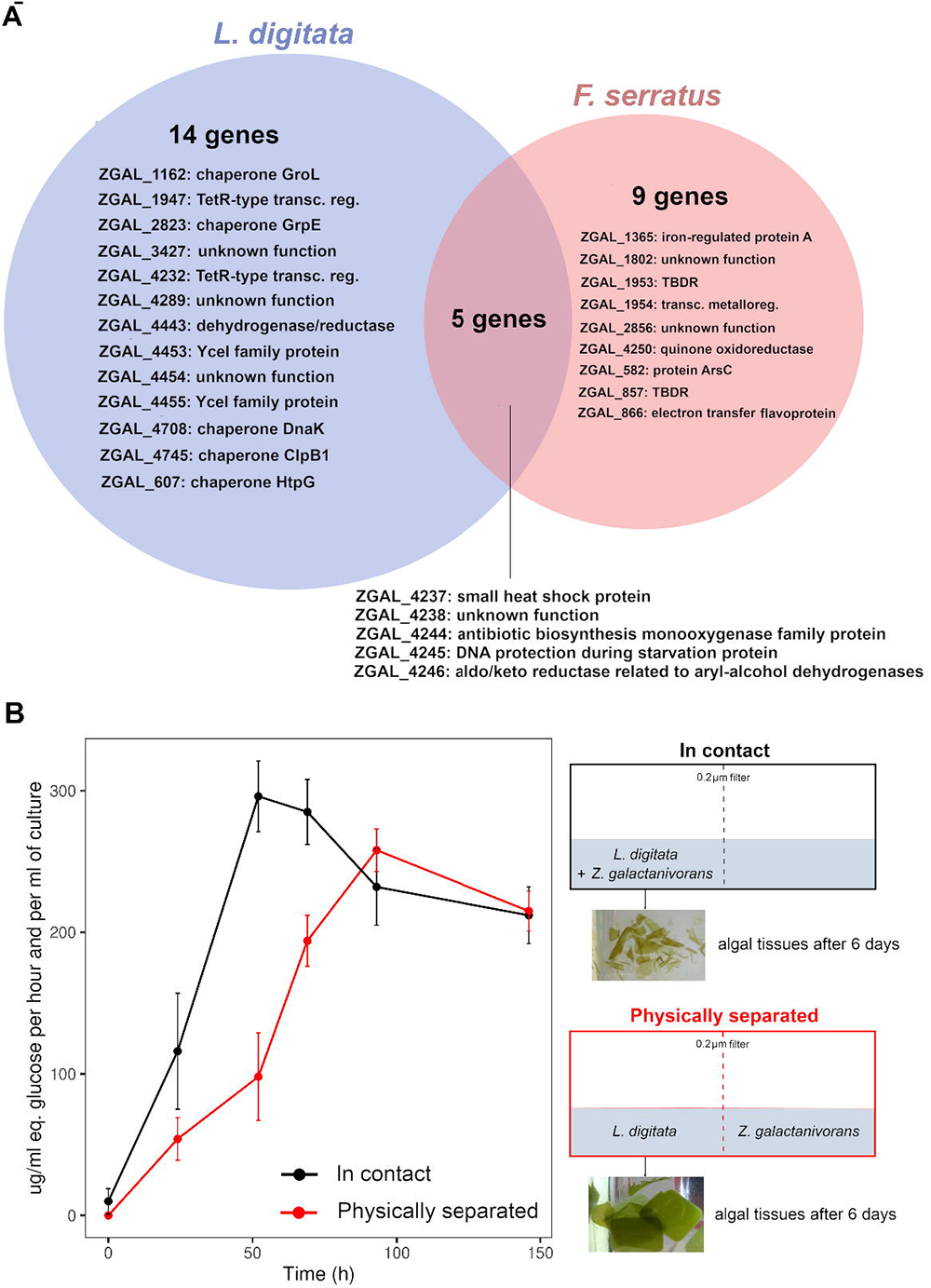
Effect of the attachment to macroalgae during the degradation. (A) Number of genes (Bonferroni-adjusted p-value < 0.05) up-regulated in algae-attached bacteria compared to free-living bacteria. The annotation of each gene is provided. (B) Alginolytic activity of the enzymes secreted when *Z. galactanivorans* was grown in contact with *L. digitata* (black) or separated from *L. digitata* by a 0.2 μm filter (red). The activity was measured in each compartment (left and right) and summed. Values are mean ± s.d. (n = 3).

To assess if algal degradation requires biofilm formation, *Z. galactanivorans* was grown either in contact or physically separated from algal pieces (**Figure 5B**). After 6 days, algal tissues were visually starting to decompose when bacteria were separated from algae, although to a lesser extent compared to the “contact” condition. Furthermore, extracellular alginolytic activity increased even without physical bacteria/algae contact and reached similar levels to that observed in the “contact” condition after 90 h.

### Comparative physiology and genomics of fresh macroalga degradation by *Zobellia*

The degrading abilities of other members of the genus *Zobellia* were investigated (**Figure 6A**). All tested *Zobellia* strains used fresh *L. digitata* tissues for their growth. *Z. galactanivorans* Dsij^T^ had the highest final cell density (OD_600_ = 1.5) and shortest generation time (5.09 h). *Z. nedashkovskayae* Asnod3-E08-A formed cell aggregates that biased OD_600_ readings, likely explaining the apparent limited growth (final OD_600_ = 0.4) and long generation time (t_gen_ = 16.33 h). Other strains showed intermediate behaviors (OD_600_ ≈ 1, 5.92 < t_gen_ < 11.74 h). These growth differences were reflected in the final aspect of macroalgal pieces. Only *Z. galactanivorans* Dsij^T^ completely broke down algal tissues after 91 h. Both *Z. nedashkovskayae* strains caused limited algal peeling and breakdown at the corners of the pieces., No visible trace of degradation was detected for other strains. A strong negative correlation was found between the number of GHs and the generation time (Spearman, rho = - 0.90; P = 0.006) (**Figure 6A, SuppTable 5**). Twenty-two out of the 305 genes up-regulated by *Z. galactanivorans* Dsij^T^ with *L. digitata* compared to maltose had no homologs in the genome of the seven other *Zobellia* strains (**SuppTable 6**). They include two GHs, *zgal_3349* (GH20 in PUL33) and *zgal_3470* (GHnc in PUL35), and a susCD-like pair (*zgal_3440, 3441)* in PUL34. Other up-regulated genes within the FCSP PUL34/35 are not conserved in all *Zobellia* strains (**Figure 6B**). Likewise, several alginolytic genes were not conserved across the genus, especially in the two *Z. roscoffensis* strains that lack 7 of them. *zgal_1182* and *zgal_4327,* encoding the extracellular endo-alginate lyases AlyA1 and AlyA7 respectively, were not conserved in the other strains *(zgal_4327)* or only found in the *Z. nedashkovskayae* strains *(zgal_1182).* Two other genes related to carbohydrate assimilation *(zgal_334* and *zgal_2296* encoding a GHnc and a lipoprotein with CBM22, respectively) are missing in five strains (**SuppTable 6**). *zgal_334* neighbors genes encoding sulfatases, fucosidases and PLs and might belong to a FCSP-targeting cluster (absent from the 51 identified PULs as the pair *susCD*-like is absent).

**Figure 6.**
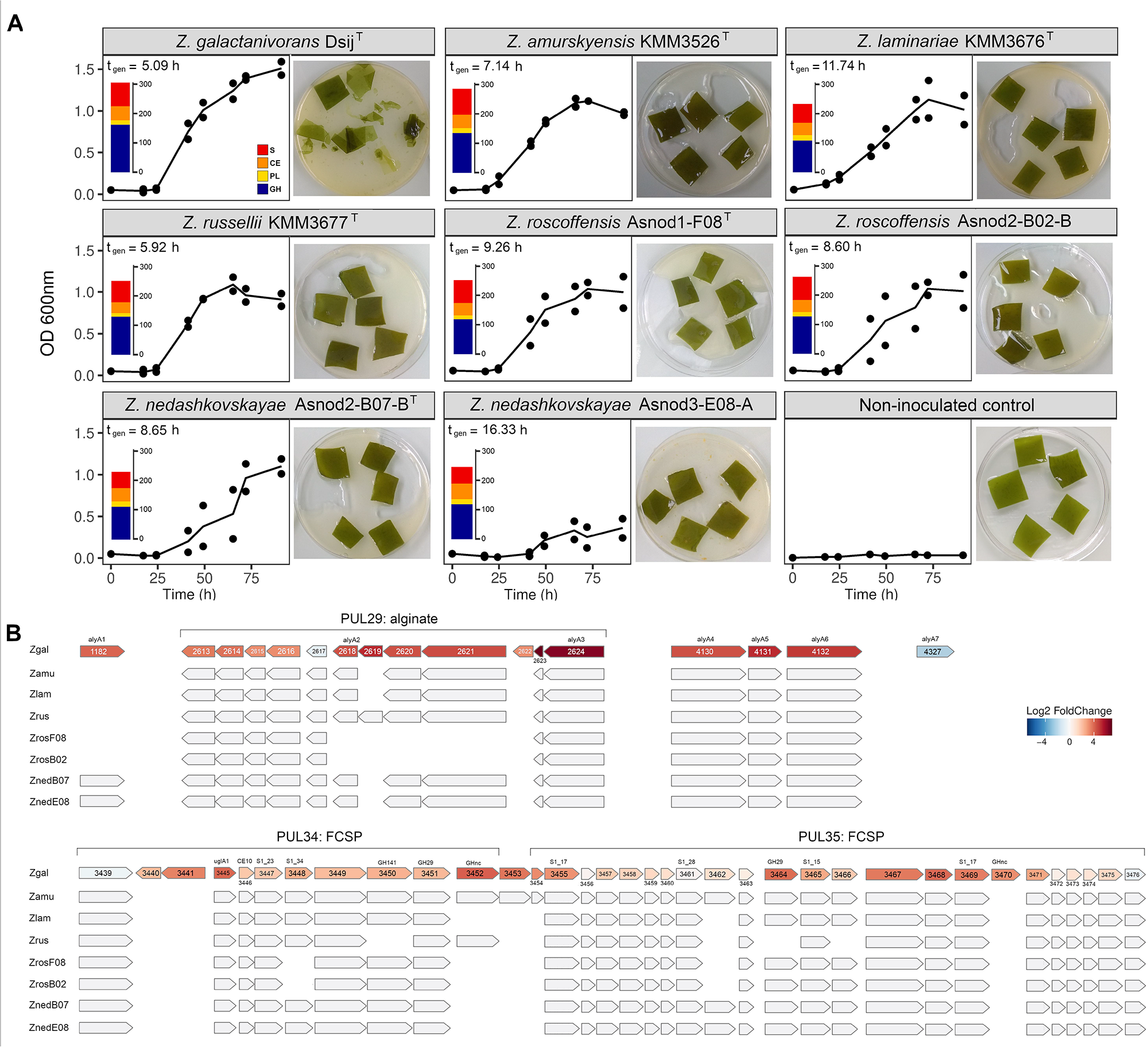
Ability of *Zobellia* spp. to use fresh *L. digitata* for its growth. (A) Growth of eight *Zobellia* strains with *L. digitata* pieces (meristem of adult individuals). The generation time tgen is indicated for each strain as well as the number of glycoside hydrolases (GH, blue), polysaccharides lyases (PL, yellow), carbohydrate esterases (CE, orange) and sulfatases (S, red) predicted in their genome (dbCAN search on the MaGe platform). Individual points for duplicate experiments are shown. Lines are means of independent replicates (n = 2 or n = 3). (B) Comparison of genomic loci among the eight *Zobellia* strains. For *Z. galactanivorans*, genes were colored according to their expression log2FC for the comparison *L. digitata* vs. maltose. Gene ID is indicated inside arrows and CAZymes and sulfatases are specified above. Top: genes involved in the alginate-utilization system. Bottom: genes contained in putative FCSP PUL34 and 35. Zgal: *Z. galactanivorans* Dsij^T^; Zamu: *Z. amurskyensis* KMM 3526^T^; Zlam: *Z. laminariae* KMM 3676^T^; Zrus: *Z. russellii* KMM 3677^T^; ZrosF08: *Z. roscoffensis* Asnod1-F08^T^; ZrosB02: *Z. roscoffensis* Asnod2-B02-B; ZnedB07: *Z. nedashkovskayae* Asnod2-B07-B^T^; ZnedE08: *Z. nedashkovskayae* Asnod3-E08-A.

## Discussion

### *Zobellia galactanivorans* Dsij^T^ acts as a sharing pioneer in brown macroalgae degradation

By degrading macroalgae, marine heterotrophic bacteria are central to nutrient cycling in coastal habitats. The ecological strategies of different functional guilds not equally equipped to process biomass were recently conceptualized (19,30,31). First, pioneer bacteria degrade complex organic matter by producing specific hydrolytic enzymes. The hydrolysate can then fuel other bacteria called exploiters or scavengers, which cannot feed on intact substrates. Such cooperative interactions were previously characterized during alginate (74) or chitin (75,76) assimilation. Hence, in nature pioneer bacteria likely control the initial attack on fresh macroalgae, a hitherto rarely studied process that cannot be fully deciphered when using purified polysaccharides or crushed algae. Here, we showed that *Z. galactanivorans* Dsij^T^ uses healthy brown algal tissues for its growth, highlighting its pioneer role in algal biomass recycling. Similar growth rates were observed with three brown algal species, and *Z. galactanivorans* completely broke down *L. digitata* tissues. Transcriptomes obtained with *L. digitata* were closest to that with alginate and FCSP, suggesting a greater capacity to access and digest ECM polysaccharides within the *L. digitata* tissues compared to *A. nodosum* and *F. serratus.* The limited degradation of Fucales tissues might originate from their higher phlorotannin content (77), possibly inhibiting CAZymes (78). In addition, *A. nodosum* induced a wider cellular response with many specific regulations. This might partly be due to the growth of antibiotic-resistant epiphytic bacteria that could have affected *Z. galactanivorans* behavior or to its much thicker and rigid thallus. Furthermore, *A. nodosum* is associated with various symbionts, especially the obligate endophytic fungus *Mycophycias ascophylli* (79) that secretes compounds potentially preventing tissue grazing and/or offering new substrate niches.

We showed that although *Z. galactanivorans* can colonize *L. digitata,* it does not require a physical contact to initiate degradation. Furthermore, only few upregulated genes were detected in surface-attached vs. free-living cells. While difficulties to extract RNA from algae-attached bacteria resulted in poor sequencing coverage, it should still have been possible to detect strong upregulations in attached cells. Overall, our results indicate that surface attachment is not required for the utilization of algal biomass by *Z. galactanivorans* Dsij^T^. This suggests a crucial role for secreted enzymes to initiate degradation, in line with the measured extracellular alginolytic activity. Constitutively expressed extracellular enzymes, such as the alginate lyases AlyA1 and AlyA7 (44), would rapidly release diffusible degradation products, allowing remote substrate sensing. We previously showed that *Z. galactanivorans* accumulates low molecular weight (LMW) alginate oligosaccharides when grown with purified alginate and algal tissues (44,55). Our results therefore confirms that *Z. galactanivorans* would be a “sharing” pioneer providing degradation products as public goods to other taxa (55), contrary to “selfish” pioneers which sequester LMW products by producing essentially surface-associated hydrolytic enzymes with minor loss of hydrolysate to the medium (80,81).

We further evidenced that this pioneer behavior can be strain-specific within the alga-associated genus *Zobellia.* All *Zobellia* spp. tested successfully grew with fresh *L. digitata* but without causing pronounced tissues damages as observed with *Z. galactanivorans*. Their catabolic profiles (**SuppTable 1**) indicate different growth capacities with purified brown algal sugars. For example, *Z. roscoffensis* strains and *Z. laminariae* KMM 3676^T^ display limited or no abilities to use alginate, FCSPs and laminarin for their growth. Hence, with macroalgae, they likely did not use these complex polysaccharides but rather fed on soluble algal exudates (e.g. mannitol). Comparative genomics suggested that CAZyme content influences the strain capacity to use and break down fresh algal tissues. In particular, some strains lack homologs of overexpressed genes contained in *L. digitata-induced* PULs targeting alginate or FCSPs. For example, *alyAl* homologs were only found in the two other strains that caused visible algal damage (Z. *nedashkovskayae* Asnod2-B07-B^T^ and Asnod3-E08-A). Accordingly, *alyA1* is known to have a crucial role in initiating algae breakdown (55). Such genes would therefore represent potential genetic determinants of pioneer bacteria.

### Deciphering the metabolic mechanisms involved in fresh tissue breakdown, including new catabolic pathways

Regardless of the algal species, the well-characterized alginolytic PUL29 was the most induced among all PULs. Alginate is the most abundant polysaccharide in brown algal ECM and likely the most accessible as it embeds the cellulose-FCSP network (11). This PUL was particularly triggered with *L. digitata,* likely reflecting the higher alginate content in this species (2) and/or an easier substrate accessibility. Furthermore, several uncharacterized PULs were triggered with macroalgae. Three out of the seven predicted FCSP PULs were significantly upregulated with macroalgae but not with extracted *A. nodosum* FCSPs, and to various degrees depending on algal species. In addition, two PULs with unclear function (PUL26 and 27) were induced with both FCSPs and macroalgae. This suggests different substrate specificities, consistent with the large structural diversity of FCSPs and crosslinkage to other compounds (4,7,82) which might not be equally extracted during purification. By preserving the original polysaccharide structure and environment, the study of fresh macroalga degradation may therefore be a more effective way to reveal specific genes crucial for macroalgae breakdown by pioneer bacteria but undetectable when using purified polysaccharides.

By contrast to alginate- and FCSP-targeting PULs, the characterized laminarin PUL11 and PUL28 were poorly regulated with the three algae. An uncharacterized β-glucan PUL4 was significantly induced only with *A. nodosum,* and also found triggered with purified laminarin in a previous study (24). As raised above, the presence of endosymbionts in *A. nodosum* could result in specific laminarin structures that might be targeted by PUL4. The absence of induction of typical laminarin PULs with macroalgae might also indicate that *Z. galactanivorans* Dsij^T^ first uses ECM polysaccharides and later access intracellular storage polysaccharides. Such a prioritization of multiple substrates within algal material was previously observed for *Bacillus weihaiensis* Alg07^T^ grown on algal powder (23). Koch *et al.* (26) showed that *Alteromonas macleodii* 83-1 prioritized laminarin over alginate and pectin when grown on a mixture of purified polysaccharides. Thus, prioritization might differ between bacterial strains and whether substrates are under soluble form or within algal tissues, underlining the importance to consider intact macroalgae to understand the pioneer behavior. Furthermore, future time-resolved transcriptome analyses could inform on regulations at different degradation stages and help decipher prioritization effects.

Besides carbohydrate utilization, our approach unveiled several traits specifically induced upon macroalgal degradation and potentially linked to the pioneer behavior, including the resistance to algal defense and T9SS. One of the algal defense mechanisms is the production of reactive oxygen species (ROS), which in *L. digitata* is partly induced by endogenous elicitors (i.e. oligo-alginates) derived from the degradation of their own cell wall (83). Breakdown of *L. digitata* tissues by *Z. galactanivorans* likely produced large amounts of elicitors and consequently triggered a massive oxidative burst, in line with the strong induction of genes encoding ROS-detoxifying enzymes in this condition. In contrast, *A. nodosum* and *F. serratus* do not respond to the addition of endogenous elicitors (84), potentially explaining the lower induction of antioxidant pathways in *Z. galactanivorans* Dsij^T^ with these algae. Another algal defense response is the emission of halogenated compounds. One vanadium-dependent iodoperoxidase (vIPO3) and a haloacid dehalogenase (HAD, (54)) were significantly up-regulated with *A. nodosum* compared with alginate and maltose respectively. HAD expression was also 3-fold higher with *L. digitata* and *F. serratus* compared to maltose, although large variations precluded significance. The induction of stress resistance mechanisms was even more pronounced in bacteria attached to *L. digitata* tissues through the expression of chaperones. Overall, our results suggest that pioneer bacteria might have evolved to cope with increasing stress levels upon algal degradation. By metabolizing toxic compounds, they might favor the growth of less stress-resistant scavenger bacteria, a hitherto overlooked additional benefit besides the opening of new substrate niches.

Specific to *Bacteroidetes,* T9SS is involved in biofilm formation, protease virulence factors delivery and secretion of polysaccharidases and cell-surface gliding motility adhesins (85,86). Here, we showed that growth with macroalgae strongly induced genes encoding T9SS components, T9SS-translocated proteins and several glycosyl transferases from families GT2 and GT4. Glycosyltransferases with a GT4_CapM-like domain were recently shown to N-glycosylate CTD in *Cytophaga hutchinsonii,* an essential step for the recognition of cargo proteins by T9SS (87). Hence, our data suggest T9SS might be a key determinant of pioneer behavior in the *Bacteroidetes* phylum, to secrete ECM-targeting CAZymes and/or attach to macroalgal surfaces.

## Conclusion

This study provides the first insights into the metabolic strategies of sharing pioneer bacteria during fresh macroalgae utilization and represents a source of potential genetic determinants for further functional characterization. Altogether, our results raised the relevance to consider the whole complexity of macroalgae tissues in further degradation studies, as it would take a step forward in the understanding of the algal biomass recycling through the identification of new metabolic pathways or the characterization of bacterial cooperative interactions.

## Supporting information

SuppTable 3

SuppTable 4

SuppTable 6

## Acknowledgments

The authors thank Tatiana Rochat for advice during transcriptomic analyses, Sébastien Colin for guidance in confocal manipulation, Philippe Potin and Cécile Hervé for helpful discussions and Yan Jaszczyszyn from the I2BC sequencing platform. This work has benefited from the facilities of the Genomer platform and from the computational resources of the ABiMS bioinformatics platform (FR 2424, CNRS-Sorbonne Université, Roscoff), which are part of the Biogenouest core facility network. This work was funded by the French Government via the National Research Agency programs ALGAVOR (ANR-18-CE02-0001-01) and IDEALG (ANR-10-BTBR-04).

## Competing interests?

The authors have no conflict of interest to declare.

## Author contributions

(according to CRediT taxonomy)

## Conceptualization

MB, TB and FT. Data curation: MB, FT. Formal analysis: MB, FT. Funding acquisition: FT. Investigation: all authors. Supervision: TB, FT. Visualization: MB. Writing original draft: MB. Writing review and editing: MB, TB and FT.

## Notes

### Competing Interest Statement

The authors have declared no competing interest.

