## Supplementary Material for "Metabolic strategies of sharing pioneer bacteria mediating fresh macroalgae breakdown"

*Corresponding author

**SUPPLEMENTARY MATERIAL**

**Supplementary Methods**

**Strains**

Eight *Zobellia* strains were used in this study, *Z. galactanivorans* Dsij^T^, *Z. amurskyensis* KMM 3526^T^, *Z. laminariae* KMM 3676^T^, *Z. russellii* KMM 3677^T^, *Z. roscoffensis* Asnod1-F08^T^, *Z. roscoffensis* Asnod2-B02-B, *Z. nedashkovskayae* Asnod2-B07-B^T^ and *Z. nedashkovskayae* Asnod3-E08-A (SuppTable 1). These strains were first grown from glycerol stock in Zobell 2216 medium at room temperature before being inoculated in minimum marine medium (24.7 g.l^-1^ NaCl, 3.08 g.l^-1^ anhydrous MgSO_4_, 4.6 g.l^-1^ MgCl_2_·H_2_O, 2 g.l^-1^ NH_4_Cl, 0.7 g.l^-1^ KCl, 0.6 g.l^-1^ CaCl_2_, 0.2 g.l^-1^ NaHCO_3_, 0.6 g.l^-1^ K_2_HPO_4_, 47.6 mM Tris-HCl pH8, 0.02 g.l^-1^ FeSO_4_·7H_2_O and vitamins [5 µg.ml^-1^ of pyridoxine HCl and orotic acid, 1 µg.ml^-1^ nicotinic acid, thiamine HCl, riboflavine, D,L-pantothenic acid, D-biotin, folic acid, cyanocobalamine and ascorbic acid, 10 µg.ml^-1^ 4-aminobenzoic acid], and complemented with 100 µg.ml^-1^ kanamycin, 100 µg.ml^-1^ streptomycin, 50 µg.ml^-1^ neomycin and 14 µg.ml^-1^ colistin, to which all the tested *Zobellia* strains are resistant). This marine minimum medium (MMM) was amended with 4 g.l^-1^ maltose as the sole carbon source. Pre-cultures were grown at room temperature under agitation for 3 days. After centrifugation (3200 g, 10 min), pellets were washed twice in 1X saline solution (24.7 g.l^-1^ NaCl, 3.08 g.l^-1^ anhydrous MgSO_4_, 4.6 g.l^-1^ MgCl_2_·H_2_O, 0.7 g.l^-1^ KCl) to remove any carbon trace. Cells were inoculated in microcosms at OD_600_ 0.05.

**Macroalgae treatment**

Healthy *Laminaria digitata*, *Fucus serratus* and *Ascophyllum* *nodosum* were collected in May 2019 at the Bloscon site (48°43’29.982’’ N, 03°58’8.27’’ W) in Roscoff (Brittany, France) in plastic bag. Algae were rinsed with 0.2 μm filtered seawater and used immediately or alternatively stored overnight in filtered seawater in 5 l bottle at 16 °C with aeration under day/night light cycle. Prior to experiment, algal tissues were cut in small pieces (ca. 2.5-3.5 cm^2^) with a sterile scalpel and immersed in 0.1 % X-100 Triton in milli-Q water for 10 min followed by 1 % povidone-iodine in milli-Q water for 5 min to clean them from the resident epibiotic microbiome. Algae pieces were rinsed by successive baths (at least five) in excess autoclaved seawater for 2 hours to remove Triton and povidone-iodine traces as well as potential algal exudates released upon cutting.

**Microcosm set up and sample collection**

*Zobellia* strains were grown at 20 °C under agitation in MMM with macroalgae pieces as the sole carbon source. Each condition was tested in triplicate. All vessels were autoclaved.

The eight *Zobellia* strains used in this study were grown in 50 ml Falcon tubes with 10 ml of MMM and three *L. digitata* pieces cut in the meristem part (up to 15 cm from the base).

*Z. galactanivorans* was grown in 250 ml erlenmeyer flask with 50 ml of MMM and 10 brown macroalgal pieces, either young *L. digitata* (< 20 cm), *F. serratus* or *A. nodosum*. For comparison it was also grown in the same conditions using 4 g.l^-1^ maltose, alginate or FCSPs. During the exponential phase, 10 ml of culture medium and 2 algal pieces were retrieved on ice for RNA extraction of the free-living and algae-attached bacteria, respectively. On ice, 0.5 volume of cold (stored at 4 °C) killing buffer (20 mM Tris-HCl pH 7.5, 5 mM MgCl_2_, 20 mM NaN_3_) was immediately added to the liquid samples to kill the cells, stop transcription and preserve RNA from degradation. After centrifugation of liquid samples (3200 g, 10 min), cell pellets were frozen in liquid nitrogen. The collected algal pieces were washed twice in 1.5 ml of killing buffer:H_2_O (1:1) in 2 ml eppendorf tubes (for 1 piece) and tubes were immersed in liquid nitrogen. Samples were stored at -80 °C before further RNA extraction. For enzymatic assays, culture medium was sampled, filtered onto a 0.2 μm syringe filter and stored overnight at 4 °C until incubation with the different polysaccharides.

Co-culture systems were used to test the growth of *Z. galactanivorans* when cultivated in contact or physically separated from the algal tissues. The systems consisted of two 100 ml reaction vessels with round bottom and a 65 mm flat edge opening (ref 0861050, Witeg, Germany), separated by a 0.2 μm mixed cellulose ester filter (ref GSWP09000, Merck, Darmstadt, Germany). Thirty milliliters of MMM were poured in the two compartments and 10 *L. digitata* pieces were immersed in one. For enzymatic assays, culture medium was sampled at different times, filtered onto a 0.2 μm syringe filter and directly incubated with alginate.

### **RNA extraction and sequencing**

Free-living bacterial cells were lysed by adding 400 μl of lysis buffer (4 M guanidine thiocyanate, 25 mM sodium acetate pH 5.2, 5 g.l^-1^ N-laurylsarcosinate) and 500 μl of phenol pH 4 and incubated 5 min at 65 °C (tubes were flipped every minute). After the addition of 500 μl of chloroform:isoamyl alcohol (IAA) (24:1), tubes were vortexed and centrifuged (16000 g, 10 min, 4 °C). The aqueous phase was recovered carefully and extraction was repeated with 250 μl of phenol and 250 μl of chloroform:IAA. Once again, the aqueous phase was recovered and 0.25 volume of 96 % molecular biology grade ethanol was added drop by drop under agitation to precipitate potential polysaccharide traces before the addition of 250 μl of phenol and 250 µl of chloroform:IAA. The final aqueous phase was recovered after vortex and centrifugation and was mixed with 500 μl of chloroform:IAA. After vortex and centrifugation (16000 g, 10 min, 4 °C), the aqueous phase was mixed with sodium acetate pH 5.5 (1:10, sodium acetate:sample) and 2.5 volumes (from the recovered sample) of 96 % molecular biology grade ethanol were added. The solution was incubated at room temperature for 2 h to precipitate RNA and centrifuged (18000 g, 45 min, 4 °C). Pellets were washed with 500 μl of 70 % molecular biology grade ethanol, centrifuged (18000 g, 20 min, 4 °C) and air-dried on ice for 1 h under chemical hood before being resuspended in 30 μl of RNAse-free water. To remove DNA contamination, Turbo DNase buffer was added (1X final) and extracts were treated 1 h at 37 °C with 2 units of Turbo DNAse (ThermoFisher, Scientific, Waltham, MA, USA). RNA was then purified on mini-columns NucleoSpin RNA Clean-up (Macherey-Nagel, Hoerdt, France) following the manufacturer's instructions. RNA was eluted in 50 μl of nuclease-free water and stored at -20 °C.

RNA from algae-attached bacteria was extracted as follows. Two algal pieces were immersed in 2 ml killing buffer:H2O (1:1) in 15 ml Falcon tubes, vortexed 30 sec and placed 7 min in an ultrasonic bath (Ultrasonic Cleaner, 45kHz, VWR) at room temperature to detach bacteria from the algal surface. Algae were removed using sterile tweezers, buffer was centrifuged (12800 g, 5 min, 4 °C) and cell pellets resuspended in 400 µl lysis buffer I. RNA extraction and DNAse treatment were then performed as described above for the free-living bacteria. To avoid RNA loss on purification columns, DNAse was inactivated by adding 0.2 volume of DNAse inactivation reagent (ThermoFisher, Scientific). Tubes were vortexed, incubated 5 min at room temperature, centrifuged (10 000 rpm, 1.5 min) and supernatant was recovered in new tubes. RNA was stored at -20 °C. The absence of DNA contamination in all RNA samples was checked by PCR with universal primers S-D-Bact- 0341-b-S-17 and S-D- Bact-0785-a-A-21 targeting the 16S rRNA gene (57). One microliter of sample was added to the PCR mixture containing 1 µl of each 5 µM primer, 10 µl of Taq 2X Master Mix (New England BioLabs, Ipswich, MA, USA) and 7 µl of nuclease-free water. The amplification program consisted of an initial step at 95 °C for 2 min followed by 30 cycles of 95 °C for 30 s, 53 °C for 30 s and 72 °C for 30 s and then hold at 72 °C for 5 min. RNA was quantified on a Qubit fluorometer (ThermoFisher Scientific) using the Qubit RNA HS assay kit and its integrity assessed on a Bioanalyzer 2100 system (Agilent Technology, Santa Clara, CA, USA) with the Agilent RNA 6000 Pico assay kit.

Paired-end RNA sequencing (RNA-seq) was performed by the Plateforme de Séquençage I2BC (UMR9198, CNRS, Gif-sur-Yvette) on a NextSeq instrument (Illumina, San Diego, CA, USA) using the NextSeq 500/550 High Output Kit v2 (75 cycles) which included a Ribo-Zero ribosomal RNA depletion step (as Illumina did no longer supply the Illumina RiboZero rRNA Removal Kit for Bacteria at that time, rRNA depletion was carried out by mixing probes from the kit adapted for bacteria to the ones adapted for mammals).

### **RNA-seq analysis**

Demultiplexed and adapter-trimmed reads were processed with the Galaxy platform (<https://galaxy.sb-roscoff.fr>). Trimmomatic (Galaxy Version 0.38.0) was used to perform quality filtering of the reads (SLIDINGWINDOW:4:20 LEADING:3 TRAILING:3 AVGQUAL:25 MINLEN:20). Transcript quantification was done using the pseudo-mapper Salmon v0.8.2 with the *Z. galactanivorans* Dsij^T^ reference genome retrieved from the MicroScope platform (<https://mage.genoscope.cns.fr/>) “zobellia_gal_DsiJT_v2”; Refseq NC_015844.1 and with the following parameters: Type of index: quasi, Perfect Hash: False, Suffix Array:1 and the default advanced parameters. Raw counts for individual samples were merged into a single expression matrix for downstream analysis. rRNA content was assessed using SortMeRNA (Galaxy Version 2.1b.4) with default parameters.
All downstream analyses were conducted in R v3.6.2. Principal Component Analysis was performed using the *DESeq2* v1.26.0 package after rlog transformation. Differential abundance analyses were also carried out with this package and genes displaying a log2 fold change |log2FC| > 2 and a Bonferroni-adjusted p-value < 0.05 were considered to be significantly differentially expressed. The upset plot was done using the *ComplexUpset* package. Hierarchical clustering was performed using the Ward’s minimum variance method. All graphics were made with the *ggplot2*.

### **Enzymatic activity assays**

One volume of 0.2 μm filtered supernatant was incubated with 9 volumes of 0.2 % macroalgal polysaccharides (alginate, agar, kappa- or iota-carrageenans) at 28 °C overnight to assess extracellular polysaccharidases activity. Incubations were performed in parallel with inactive enzymes (boiled 15 min at 98 °C) as control. The amount of reducing ends released in the incubation mix was quantified using the ferricyanide assay as follows. Twenty microliters of 5X ferricyanide (1.5 g potassium hexacyanoferrate III, 140 g dehydrated sodium carbonate, 25 mM NaOH) was added to 80 μl of the incubation mixtures in 96-wells microplate. A standard curve of glucose was prepared by mixing 20 μl of 5X ferricyanide with 70 μl of milliQ-water and 10 μl of glucose at different concentrations (0.25–1.5 mM). Plate was agitated 30 s at 1000 rpm, incubated 10 min at 98 °C then cooled down 10 min at 4 °C. Each sample was assayed in technical duplicates. Eighty-five microliters of each well were transferred in a 96-wells microplate and the absorbance was measured at 420 nm using microplate reader (Spark Tecan). The amount of reducing-ends was estimated in glucose equivalent using the glucose standard curve. For each sample, the value measured with the boiled enzymes was subtracted. Finally, the mean value (n=3) measured for the non-inoculated microcosms was subtracted. Significant differences (P<0.05) from 0 were tested using t-tests.

### **CARD-FISH**

Medium supernatant was collected after 65 h from the microcosm experiment implemented to perform the transcriptome analysis. To assess algal surface colonization, CARD-FISH assays were performed on samples collected at different time points of an independent microcosm experiment with *L. digitata* tissues as the carbon source. Supernatants and algal pieces were fixed overnight at 4 °C with 2 % paraformaldehyde, washed twice in PBS and stored in PBS:EtOH (1:1 v/v) at -20 °C.
Algal pieces were peeled using a sterile scalpel. Small portions were cut, dipped in 0.1% ME agarose, placed on a glass slide and air-dried for > 30 min at 37 °C to attach the tissues on the slide. Slides were then dipped 10 min in ethanol with increasing concentration (50, 80 and 96 %). Cells were permeabilized by incubating the slides in lysozyme (10 mg.ml^-1^ in 0.05 M EDTA, 0.1 M Tris-HCl pH8 in milli-Q water, 15 min, 37 °C) in Petri dishes. After being washed twice in 50 ml of milli-Q water, slides were incubated 30 min in 0.15 % H_2_O_2_ in methanol to inactivate endogenous peroxidases, washed 5 min in 50 ml of 1X PBS and shortly in 50 ml of milli‑Q water. Algal samples were air-dried before being covered by the preheated hybridization mix (0.9 M NaCl, 20 mM Tris-HCl pH8, 35 % formamide, 1 % blocking reagent, 10 % dextran sulfate, 0.02 % SDS, 28 nM ZOB137 probe and each helper). Slides were placed in Petri dishes separated by spacers in a preheated humid chamber containing tissue paper soaked with 2 ml of a solution containing 35 % formamide and 0.9 M NaCl in milli-Q water and incubated for 2.5 h at 46 °C. Slides were first washed briefly in preheated washing buffer (70 mM NaCl, 1 M Tris-HCl pH8, 5 mM EDTA, 0.01 % SDS) to remove the excess of working solution followed by a 15 min incubation in the same buffer at 48 °C and 15 min in 1X PBS. They were air-dried and, as described for the hybridization step, algal samples were covered with a detection solution (1X PBS, 2 M NaCl, 0.1 % blocking reagent, 10 % dextran sulfate, 0.0015 % H_2_O_2_ and 1 μg.ml^-1^ of Alexa546 labeled tyramides) and incubated 45 min in humid chamber at 46 °C for amplification. Slides were washed 10 min in 1X PBS in the dark, twice briefly in 50 ml of milli-Q water and dehydrated in 50 ml of 96% ethanol for 1 min. Algal samples were covered with DAPI (1 μg.mg^-1^) for 10-15 min in the dark then washed briefly twice in milli-Q water and in 96 % ethanol. Slides were mounted once samples were completely dry with Citifluor:Vectashield (3:1). Cells on algal tissues were visualized with a confocal microscope Leica TCS SP8 equipped with HC PL APO 63x/1.4 oil objective using the 488 and 638 nm lasers to detect Alexa488 signal and algal autofluorescence signal, respectively. Z-stack images were collected (layers of 0.29 μm thickness) by using 1024x1024 scan format and a scan speed of 400 Hz. Z-stacks were visualized using the surface channel mode of the 3D viewer module implemented in the Leica Las X software.

Free-living bacteria were harvested using a vacuum pump onto a 0.2 μm Whatman polycarbonate membrane placed on a 0.45 cellulose acetate membrane. Membranes portions were processed as described for the algal samples (without slide-attachment at the beginning of the protocol). Endogenous peroxidases inactivation was done in 0.1 M HCl (1 min) followed by 10 min in freshly prepared 3 % H_2_O_2._ Membrane portions were visualized with a Leica DMi8 epifluorescent microscope equipped with an oil objective 63X and a Leica DFC3000 G camera (Wetzlar, Germany).

**Supplementary Tables**

SuppTable 1: Ability of the eight *Zobellia* strains tested in this study to use purified brown algal polysaccharides. NA, no data available.

|  | **Alginate** | **FCSP** | **Laminarin** | **Mannitol** | **Reference** |
| --- | --- | --- | --- | --- | --- |
| *Z. galactanivorans* Dsij^T^ | + | + | + | + | Barbeyron et al., 2001 |
| *Z. amurskyensis* KMM 3526^T^ | ± | + | + | + | Nedashkovskaya et al., 2004 |
| *Z. laminariae* KMM 3676^T^ | ± | ± | ± | + | Nedashkovskaya et al., 2004 |
| *Z. russellii* KMM 3677^T^ | NA | NA | NA | NA | Nedashkovskaya et al., 2004 |
| *Z. roscoffensis* Asnod1-F08^T^ | - | - | - | + | Barbeyron et al., 2021 |
| *Z. roscoffensis* Asnod2-B02-B | - | - | - | + | Barbeyron et al., 2021 |
| *Z. nedashkovskayae* Asnod2-B07-B^T^ | + | + | + | + | Barbeyron et al., 2021 |
| *Z. nedashkovskayae* Asnod3-E08-A | + | + | + | + | Barbeyron et al., 2021 |

SuppTable 2: General characteristics of the different RNA samples.

| **Sample** | **Type** | **OD_600_ at sampling time** | **RNA concentration (ng/μl)** | **ratio A260/280** | **ratio A260/230** | **Genome coverage (%)** | **Prokaryotic rRNA (%)** | **Eukaryotic rRNA (%)** | **Number of fragments post trimming** | **Genes not covered (%)** |
| --- | --- | --- | --- | --- | --- | --- | --- | --- | --- | --- |
| FL_Ldig1 | Free-living | 0.3 | 2.8 | 1.7 | 0.7 | 74.7 | 6.1 | 0.7 | 3.09x10^7^ | 0.1 |
| FL_Ldig2 | Free-living | 0.4 | 19.8 | 1.8 | 1.4 | 58.6 | 6.4 | 0.7 | 2.90x10^7^ | 0 |
| FL_Ldig3 | Free-living | 0.3 | 3.9 | 1.7 | 0.7 | 92.9 | 27.0 | 1.0 | 1.09x10^7^ | 1 |
| FL_Fser2 | Free-living | 0.2 | 33.2 | 2.1 | 0.6 | 80.0 | 2.1 | 0.8 | 2.62x10^7^ | 0 |
| FL_Fser3 | Free-living | 0.3 | 21.3 | 2.1 | 0.9 | 43.5 | 1.5 | 0.7 | 1.78x10^7^ | 0.5 |
| FL_Anod1 | Free-living | 0.4 | 35.8 | 1.6 | 1.1 | 68.7 | 1.2 | 0.1 | 1.25x10^7^ | 1.7 |
| FL_Anod2 | Free-living | 0.4 | 29.0 | 1.5 | 0.9 | 78.6 | 2.7 | 1.0 | 1.24x10^7^ | 0.1 |
| FL_Anod3 | Free-living | 0.5 | 90.0 | 2.1 | 1.6 | 59.1 | 0.8 | 0.1 | 3.22x10^7^ | 0.2 |
| FL_Maltose1 | Free-living | 0.5 | 187.0 | 2.1 | 2.0 | 87.5 | 6.0 | 0.2 | 9.51x10^6^ | 0.5 |
| FL_Maltose2 | Free-living | 0.5 | 185.0 | 2.1 | 1.3 | 90.5 | 12.6 | 0.1 | 1.11x10^7^ | 0.3 |
| FL_Maltose3 | Free-living | 0.4 | 232.0 | 2.1 | 2.0 | 90.6 | 11.8 | 0.1 | 1.57x10^7^ | 0.1 |
| FL_Alginate1 | Free-living | 0.4 | 365.0 | 1.7 | 0.9 | 93.6 | 6.9 | 0.2 | 1.21x10^7^ | 0.1 |
| FL_Alginate2 | Free-living | 0.3 | 128.0 | 1.8 | 0.7 | 93.3 | 6.6 | 0.2 | 1.46x10^7^ | 0.1 |
| FL_Alginate3 | Free-living | 0.4 | 1340.0 | 1.9 | 1.1 | 94.5 | 7.8 | 0.1 | 3.31x10^7^ | 0 |
| FL_FCSP1 | Free-living | 0.2 | 117.0 | 1.6 | 0.9 | 95.6 | 4.4 | 0.0 | 3.39x10^7^ | 0 |
| FL_FCSP2 | Free-living | 0.2 | 112.0 | 1.6 | 0.8 | 95.2 | 4.1 | 0.0 | 3.10x10^7^ | 0.1 |
| FL_FCSP3 | Free-living | 0.2 | 91.7 | 1.6 | 0.9 | 95.5 | 6.4 | 0.1 | 3.24x10^7^ | 0 |
| Att_Ldig1 | Attached | 0.3 | 7.2 | 1.6 | 0.1 | 6.2 | 4.6 | 28.8 | 2.62x10^7^ | 3.5 |
| Att_Ldig2 | Attached | 0.4 | too low | 1.7 | 0.4 | not sequenced | | | | |
| Att_Ldig3 | Attached | 0.3 | too low | 1.4 | 0.1 | 24.1 | 8.5 | 56.0 | 2.63x10^7^ | 5 |
| Att_Fser2 | Attached | 0.2 | 13.0 | 1.0 | 0.1 | 69.1 | 0.5 | 3.9 | 7.96x10^6^ | 0 |
| Att_Fser3 | Attached | 0.3 | 13.7 | 0.8 | 0.2 | 32.4 | 4.0 | 16.5 | 1.54x10^7^ | 27 |
| Att_Anod1 | Attached | 0.4 | 2.4 | 0.7 | 0.2 | 15.1 | 2.8 | 70.7 | 1.53x10^7^ | 1 |
| Att_Anod2 | Attached | 0.4 | too low | 0.7 | 0.2 | 35.5 | 4.0 | 54.5 | 2.28x10^6^ | 60 |
| Att_Anod3 | Attached | 0.5 | too low | 0.7 | 0.2 | 19.7 | 3.3 | 62.1 | 7.88x10^6^ | 41 |

SuppTable 3 (Excel file): Genes upregulated either with *Laminaria digitata*, *Fucus serratus* or *Ascophyllum nodosum* when compared either with maltose, alginate or FCSP. Log2FC is indicated in red if > 0 or green if < 0 and in bold of the corresponding P-value < 0.05.

SuppTable 4 (Excel file): Genes upregulated with at least one macroalgae when compared both with the polysaccharides and maltose (Bonferroni-adjusted p-value < 0.05 and log2FC > 2). MA: Macroalgae. “L” means they are induced with *Laminaria digitata*, “F” with *Fucus serratus* and “A” with *Ascophyllum nodosum*. Genes likely involved in antioxidant activity were highlighted in grey, genes involved in T9SS in orange and GH or PL in purple. Homologous genes were searched in the genome of the other *Zobellia* spp. used in this study. Green = homolog found; Yellow = homolog not found. Zgal: *Z. galactanivorans* Dsij^T^; Zamu*: Z. amurskyensis* KMM 3526^T^; Zlam: *Z. laminariae* KMM 3676^T^; Zrus: *Z. russellii* KMM 3677^T^; ZrosF08: *Z. roscoffensis* Asnod1-F08^T^; ZrosB02: *Z.* *roscoffensis* Asnod2-B02-B; ZnedB07: *Z. nedashkovskayae* Asnod2-B07-B^T^; ZnedE08: *Z. nedashkovskayae* Asnod3-E08-A.

SuppTable 5: Spearman correlation between the number of GHs, PLs, CEs or Sulfatases with the generation time. *Z. nedashkovskayae* Asnod3-E08-A was removed from the analysis as the measured OD_600_ was not reliable given the formation of aggregates.

|  | **rho** | **p-value** |
| --- | --- | --- |
| **GH** | -0.90 | 0.006 |
| **PL** | 0.45 | 0.305 |
| **CE** | -0.20 | 0.667 |
| **Sulfatases** | -0.52 | 0.229 |
| **Total** | -0.64 | 0.139 |

SuppTable 6 (Excel file): Genes upregulated with *Laminaria digitata* when compared to maltose (Bonferroni-adjusted p-value < 0.05 and log2FC > 2). Homologous genes were searched in the genome of the other *Zobellia* spp. used in this study. Green = homolog found; Yellow = homolog not found. Zgal: *Z. galactanivorans* Dsij^T^; Zamu: *Z. amurskyensis* KMM 3526^T^; Zlam: *Z. laminariae* KMM 3676^T^; Zrus*: Z. russellii* KMM 3677^T^; ZrosF08: *Z. roscoffensis* Asnod1-F08^T^; ZrosB02: *Z. roscoffensis* Asnod2-B02-B; ZnedB07*: Z. nedashkovskayae* Asnod2-B07-B^T^; ZnedE08: *Z. nedashkovskayae* Asnod3-E08-A.

**Supplementary Figures**

SuppFigure 1: CARD-FISH assays for the *Zobellia*-specific detection in microcosms containing *Fucus serratus* or *Ascophyllum nodosum* tissues. CARD-FISH assays were performed with the *Zobellia*-specific probe ZOB137 on culture medium from *F. serratus-* and *A. nodosum*-containing microcosms that displayed a bacterial growth.

SuppFigure 2: Normalized expression of the genes contained within the PULs shown in Figure3B. *: genes significantly overexpressed in the corresponding condition compared with maltose. Blue: *L. digitata*; Red*: F. serratus*; Green: *A. nodosum*; Grey: Maltose; Yellow: Alginate; Brown: FCSPs

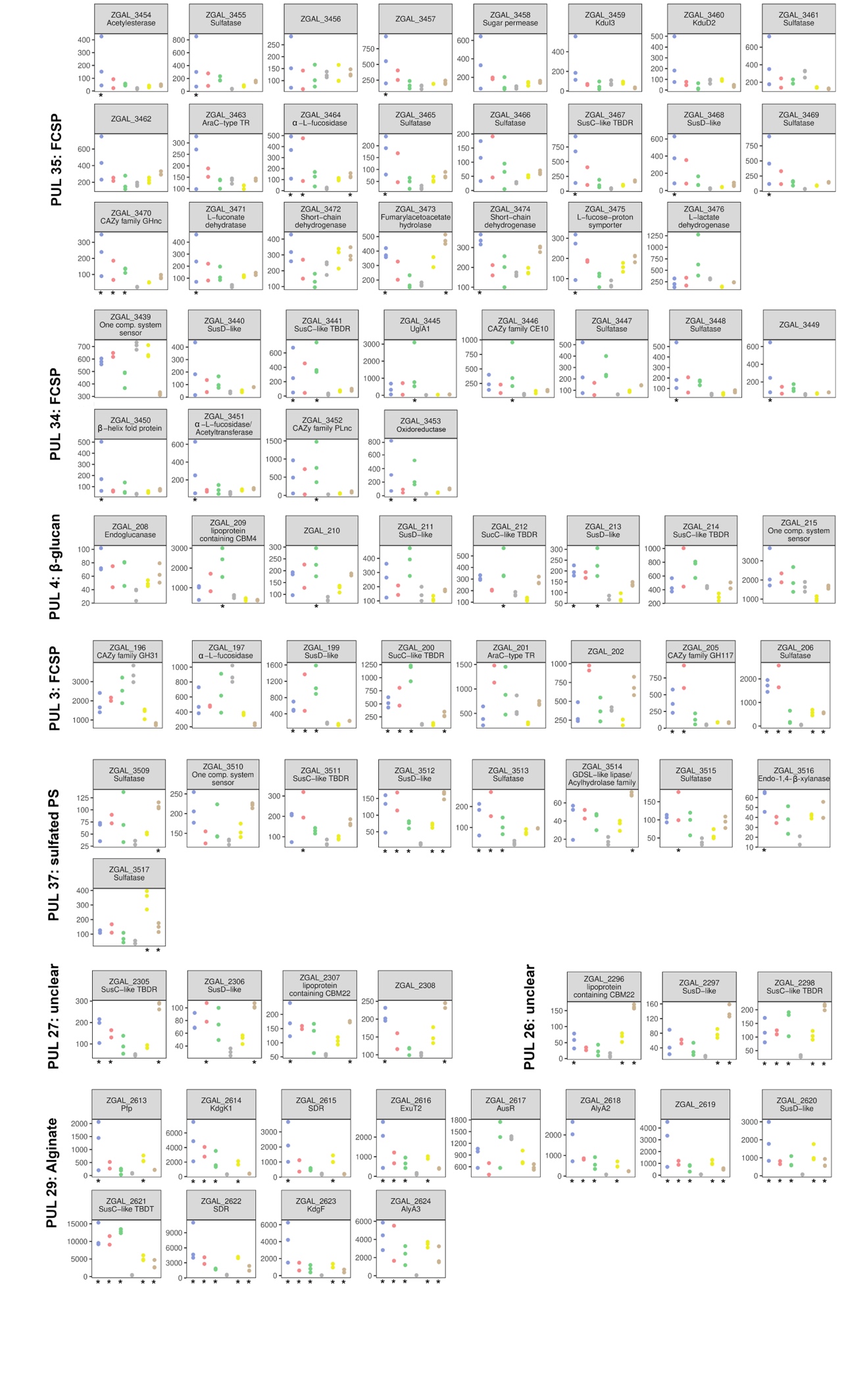
